# Genus-wide homologous recombination of tail fibers maintains tailocin diversity in *Pectobacterium*

**DOI:** 10.1101/2025.07.30.667677

**Authors:** Lakhansing A. Pardeshi, Anne Kupczok, Dick de Ridder, Sandra Smit, Theo A. J. van der Lee

## Abstract

Due to their ability to kill closely related strains, phage tail-like bacteriocins, also called tailocins, play an important role in shaping bacterial communities. One such tailocin, called carotovoricin, is also known to be present in the *Pectobacterium* genus. However, little is known about its evolutionary dynamics and the scope of impact on species interactions in this genus. To investigate the diversity and evolution of carotovoricin, we performed a genus-wide, phylogenetically-structured pangenome study. This analysis inferred that the gene cluster responsible for carotovoricin biosynthesis is conserved across the genus and is located in the same gene neighborhood in all the species. Within the carotovoricin cluster, the tail fiber genes, which determine the host range specificity, exhibit high variability and discordance with the species phylogeny. We show evidence for an evolutionary mechanism involving recombination-mediated exchange of these tail fiber loci across the entire *Pectobacterium* genus, which complements the previously known mechanism for DNA sequence inversion to maintain tailocin polymorphism at the population level. In addition, the ability to exchange tail-fiber loci in a highly targeted and genus-wide manner could influence the community dynamics in nutrient rich environments such as infected plant tissues. In conclusion, the strong signal for carotovoricin retention and ability to exchange tail fibers indicates that it significantly contributes to the community interactions of the *Pectobacterium* phytopathogens.

**Significance Statement:** A widespread presence of tailocins among various gram-negative bacteria and maintenance of their tail fiber diversity underscore their role in inter-bacterial interactions. A tailocin is also found to be conserved in *Pectobacterium*, a pathogen causing soft rot. However, the mechanism maintaining the diversity of the tailocin tail fibers, which enable recognition of the target bacteria, is not yet completely understood. Here, we characterized the genomic diversity of this tailocin and discovered that the diversity is maintained through the exchange of the tail fiber locus DNA across the genus.

## Introduction

Bacterial genomes harbor integrated temperate bacteriophages (called prophages) (Casjens 2003; Costa et al. 2018; Sharma et al. 2023; Varani et al. 2013), as well as defective prophages that degrade over time (Bobay et al. 2014). Evidence suggests that some defective prophages, lacking capsid and replication genes, were repurposed as phage tail-like bacteriocins, also known as tailocins (Borowicz, Krzyżanowska, Narajczyk, et al. 2023; Nakayama et al. 2000; Stice et al. 2023; Yamada et al. 2006). Structurally, tailocins resemble “headless phages” with tail and tail fibers forming a functional unit to kill target bacteria by puncturing the cell membrane (Scholl 2017). These tailocins are widespread throughout gram-negative bacteria (Borowicz, Krzyżanowska, Narajczyk, et al. 2023; Ghequire and De Mot 2014; Ishii et al. 2024; Kamimiya et al. 1977; G. W. Yao et al. 2017) and are divided into two classes, based on having a rigid (R-type) or flexible (F-type) core structure. The F-type tailocins are noncontractile and flexuous. In contrast, R-type tailocins have a rigid contractile tail, formed of a hollow tube encapsulated by a contractile sheath, attached to a baseplate along with tail fibers (Desfosses et al. 2019; Ge et al. 2015). The tail fibers of a contractile tailocin recognize and bind the target bacterial cell surface receptors, followed by contraction of the sheath and extension of the core tube.

As the tail fibers enable the recognition of specific target cell membrane receptors, tailocins offer an effective and targeted mechanism of killing co-occurring bacteria (Bhattacharjee et al. 2022). Normally, the target range is very narrow and limited to strains of the same species or phylogenetically close relatives (Borowicz, Krzyżanowska, Narajczyk, et al. 2023). Increasing the variability of tail fiber genes extends the recognition spectrum. Bacteria possess various mechanisms to increase the tailocin target range, such as recombination of tail fiber genes to produce new sequence combinations (Baltrus, Clark, Smith, et al. 2019), the presence of multiple tail fiber genes within tailocin loci (Dorosky, Pierson, et al. 2018), and tail-fiber locus DNA inversion by invertase (Nguyen et al. 2001). These variations in the tail fibers yield a diverse pool of tailocins that are maintained in a species (Backman, Latorre, et al. 2024).

A tailocin was discovered in the global phytopathogen *Pectobacterium carotovorum*, hence named carotovoricin (Itoh et al. 1978; Kamimiya et al. 1977). Subsequently, it was shown to be present across the entire *Pectobacterium* genus (Arizala and Arif 2019; Borowicz, Krzyżanowska, Sobolewska, et al. 2025; Pardeshi et al. 2025). Carotovoricin is produced at the basal level by most of the *Pectobacterium* species, which results in inter-species competitive interactions (Borowicz, Krzyżanowska, Sobolewska, et al. 2025). Although maintenance of diversity in tailocin tail fibers is known in other species, such as *Pseudomonas* (Backman, Burbano, et al. 2024), the existence and scope of such diversity are not yet known for the *Pectobacterium* genus. *Pectobacterium* pathogens are known to cause multi-species infections (Motyka-Pomagruk et al. 2021; Smoktunowicz et al. 2022; Zhou et al. 2022); they also show strain-specific competitive interactions, where certain strains can inhibit others, but none can inhibit the entire species (Barny et al. 2024). A systematic study of tailocin diversity at the genus level would allow understanding the broader inter-strain and inter-species interactions. It can further help to understand how carotovoricin shapes the community dynamics of co-inhabiting *Pectobacterium* strains and species.

We recently applied a comparative pangenomics approach to identify orthologous prophages and their contribution in the *Pectobacterium* genus pangenome dynamics (Pardeshi et al. 2025). Here, we elaborate our approach to systematically study the extent of conservation of the carotovoricin-biosynthesis gene cluster in the *Pectobacterium* genus. Furthermore, we study local variation within the carotovoricin cluster to understand the polymorphic nature of tailocin tail fibers. Subsequently, we evaluate the evolution of the genes within the carotovoricin cluster to help understand the underlying mechanism generating polymorphisms. Finally, we discuss the potential role of carotovoricin in the intra- and inter-species community dynamics observed in *Pectobacterium* species and the potential applications of leveraging tailocins as biocontrol strategy.

## Results

### Carotovoricin is conserved in the *Pectobacterium* genus

We recently constructed a *Pectobacterium* genus pangenome with PanTools (Jonkheer et al. 2022; Sheikhizadeh et al. 2016) and discovered the contribution of prophages to pangenome growth and the emergence of new lineages (Pardeshi et al. 2025). This pangenome comprises 454 *Pectobacterium* genomes (g_1 to g_454 in Supplementary Table S1), where 1,977,865 protein-coding genes were grouped into 30,156 homology groups. To study the dynamics of 1,319 putative prophages identified in these genomes, we represented prophages as a sequence of homology groups and traced these homology group signatures across the pangenome. We identified orthologous prophages by quantifying syntenic conservation of homology group signatures between prophages and dereplicated the prophages to 436 clusters (see Methods, Supplementary Tables S2, S3). The largest of these clusters, containing 428 orthologous prophages, was identified as the phage tail-like bacteriocin carotovoricin, based on the functional annotation of homology groups. Each carotovoricin region contained genes encoding the core structural proteins necessary for tailocin production: holin, lytic transglycosylase, tail sheath protein, tail tube protein, tail assembly protein, tail tape measure protein, baseplate proteins, tail spike protein, and tail fiber proteins (Figure 1a).

**Figure 1:**
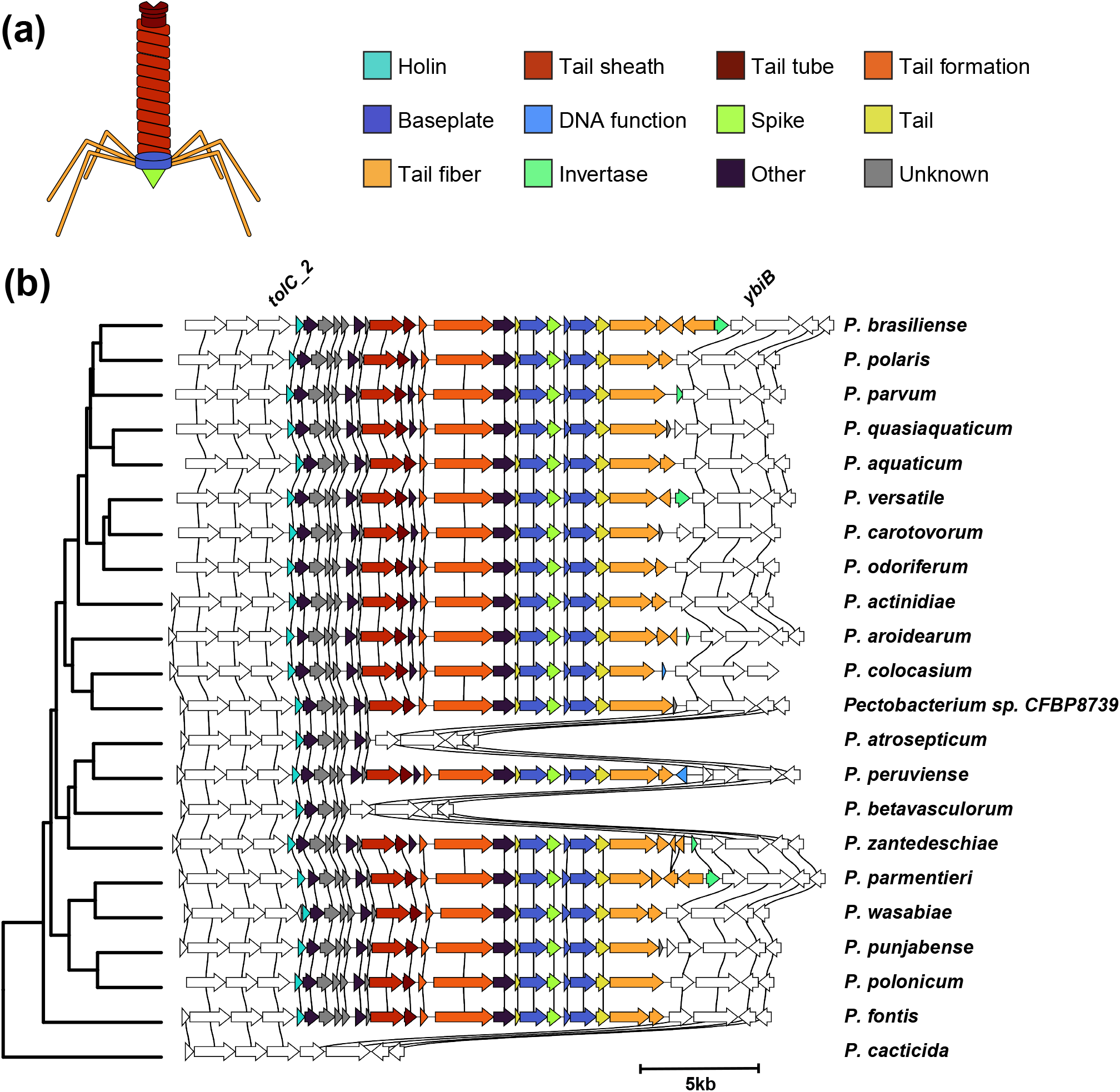
Carotovoricin cluster across the *Pectobacterium* genus. (a) An illustration of a carotovoricin particle where structural units are colored to match the carotovoricin biosynthesis cluster alignment shown in panel b. (b) Carotovoricin biosynthesis gene clusters are ordered along the phylogeny of the *Pectobacterium* genus. Genes in 5kb regions flanking the carotovoricin clusters are shown in white. Genes belonging to the same homology group are connected with vertical black lines.

We observed a single copy of the carotovoricin biosynthesis gene cluster in 428 *Pectobacterium* genomes (Supplementary Table S4). Of these, 365 were present on a single contig and the remaining were split over two contigs due to fragmented genome assemblies. Because of the presence of such fragmented assemblies in the public repository, a 5kb region flanking the carotovoricin was used to determine the flanking gene structural conservation rather than a fixed number of neighboring genes. This gene neighborhood flanking carotovoricin cluster, comprising three to four genes, is syntenically conserved across the entire *Pectobacterium* genus (Figure 1b), suggesting that a common ancestor acquired the carotovoricin cluster. Genes *tolC_2* and *ybiB* mark the upstream and downstream boundary of the carotovoricin biosynthesis gene clusters in all *Pectobacterium* species.

Complete carotovoricins encoding all core structural genes were found to be present in all but three *Pectobacterium* species (Figure 1b), detailed below. In addition to the three species-wide losses, there were seven strains from various *Pectobacterium* species that showed disruption of the carotovoricin cluster (Supplementary Results). Species with a complete carotovoricin contained 18 to 27 genes. Most of these genes were conserved, as evidenced by a single homology group per gene across all species. A partial carotovoricin cluster was observed in the *P. betavasculorum* and *P. atrosepticum* genomes, where only the first five and seven genes from the 5’ end were found to be present, respectively. The genes in these partial carotovoricins are part of the first transcriptional unit out of the four identified in the carotovoricin cluster (Yamada et al. 2006). Publicly available transcriptomics data (Kang et al. 2022) suggests that the genes within this partial carotovoricin cluster are expressed and regulated in *P. atrosepticum* (see Methods); this could not be verified for *P. betavasculorum*, due to a lack of data. Based on a phylogeny built on 1,949 core genes in the pangenome (Supplementary Figure S1), *P. betavasculorum* and *P. atrosepticum* share a common ancestor with *P. peruviense* and *P. zantedeschiae* respectively, both with intact carotovoricins. Given the difference in the deleted carotovoricin cluster and the phylogenetic distance to the closest carotovoricincarrying species, it seems that carotovoricin cluster was degraded independently in *P. betavasculorum* and *P. atrosepticum*. The third species without carotovoricin is *P. cacticida*, which lacks carotovoricin completely and shows no remnants. A recent phylogenomic analysis proposed a reclassification of *P. cacticida* into a new genus, *Alcorniella comb. nov* (Jonca et al. 2024), which likely explains the lack of carotovoricin in this genome. Overall, carotovoricin is conserved and transmitted vertically in the *Pectobacterium* genus.

### Tail fiber genes show higher variation than other carotovoricin genes

To understand the inter-species variation in carotovoricin, we investigated if it has further evolved independently in each species. To avoid analysis artifacts due to assembly quality, we selected carotovoricin cluster from 365 genomes in which it was present on a single contig together with the flanking genes. The optimal homology grouping that accounted for sequence diversity (see Methods) resulted in a single homology group for each of the carotovoricin core structural genes across the *Pectobacterium* genus. However, we found a total of 130 homology groups mapped to carotovoricin regions across the *Pectobacterium* genus, in contrast to the anticipated small set of carotovoricin-related homology groups. Closer inspection of the carotovoricin cluster across genomes showed a conserved region where the genes were represented by a single homology group, and a variable region where the genes were represented by multiple homology groups (Figure 1b, Supplementary Table S5).

First, we observed presence-absence variation of the nitric oxide deoxygenase gene (Supplementary Table S5) (present in 88/365 the genomes) (Figure 2a). The gene encoding the tail tape measure protein, located between the tail assembly chaperone and P2 GpU family protein, also showed variation across the *Pectobacterium* genus and was represented by 9 homology groups. Finally, the 3’ region flanked by the tail protein and the *ybiB* gene was represented by multiple homology group combinations across the pangenome annotation graph (Figure 2a). Interestingly, the invertase *ein*, responsible for the inversion of tail fiber locus DNA, also showed presence-absence variation: it was found in 47% (172/365) of the carotovoricins. A similar degree of diversity in carotovoricin cluster was also observed within a species, as exemplified by *P. brasiliense* and *P. versatile* species (Supplementary Figure S4a-b). In summary, the carotovoricin cluster seems to be broadly organized into two differentially conserved loci, both at the genus and species levels.

**Figure 2:**
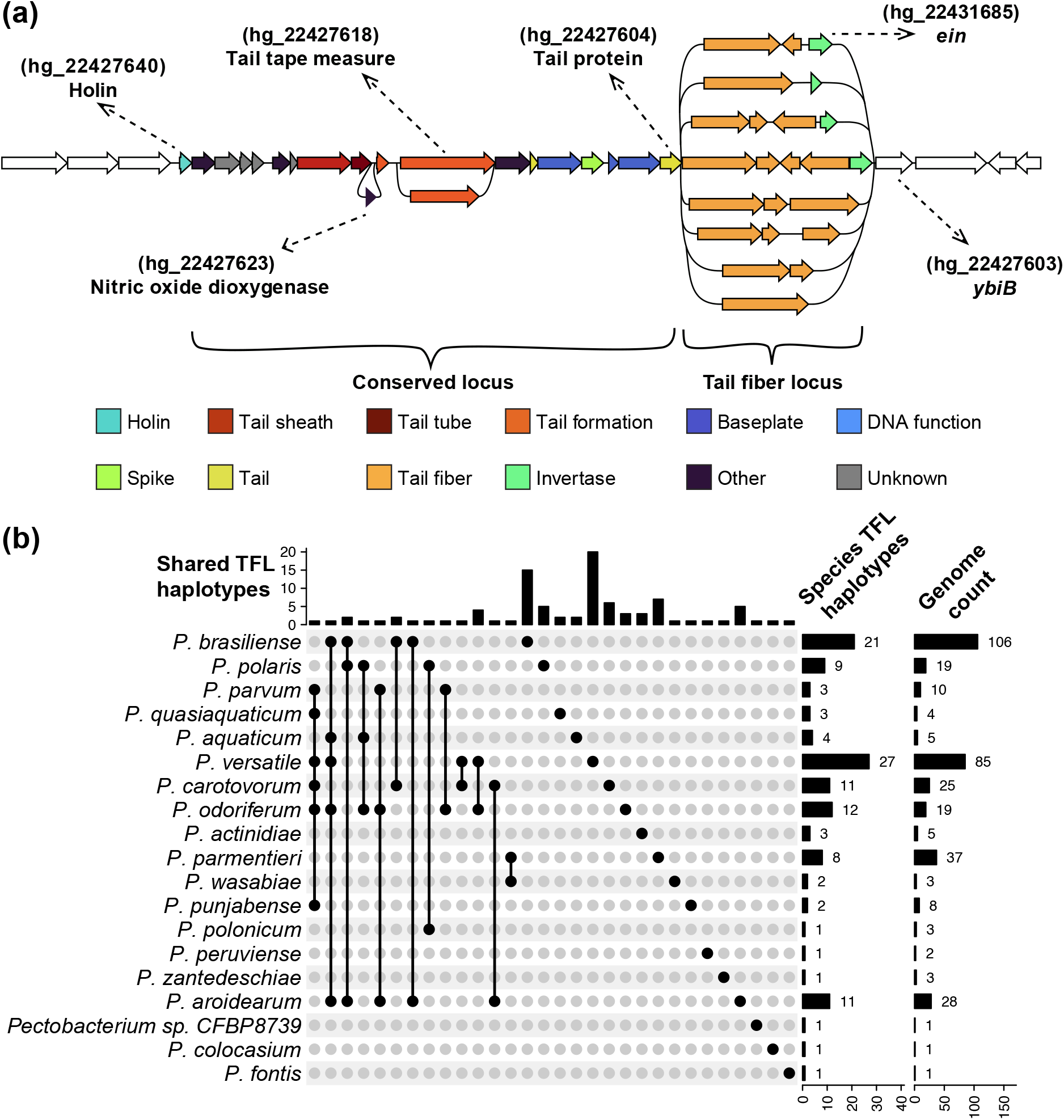
Carotovoricin locus variation. (a) An illustration of gene-level variation in carotovoricin cluster, with gene presence-absence variation shown as loops. Flanking genes are represented as white arrows. The conserved locus of carotovoricin is defined as genes encompassing holin and tail protein. The variable tail fiber locus is defined as the genes between the tail protein and *ybiB*. The unique homology group combinations in the TFLs across the *Pectobacterium* genus are used to define haplotypes. Pangenome homology group identifiers for the relevant genes are mentioned in parenthesis. (b) The UpSet plot shows the number of *Pectobacterium* species that share a TFL haplotype. Species are shown along the rows of the UpSet plot and the black dotted lines show the intersection of TFL haplotypes. The top bar chart shows number of unique TFL haplotypes shared in the respective intersection. The two bar charts on the right show total number of TFL haplotypes per species and number of genomes for each species in the pangenome, respectively.

As illustrated in Figure 2a, we define the carotovoricin cluster region spanning from the holin to the tail protein encoding gene as the conserved locus, and the remaining region as the tail fiber locus (TFL), for subsequent analysis. We further used the uniquely ordered sequence of homology groups in the TFL to define TFL haplotypes. For example, two TFLs containing 4 genes from groups A-B-D-E (in that order) are considered the same, meaning the proteins are sufficiently similar to be grouped into the same homology groups. On the other hand, A-B-D-E and A-B-E-D are considered different haplotypes. In total, there were 93 TFL haplotypes in the *Pectobacterium* genus (Supplementary Table S6). The TFL lengths ranged between 2 and 6 kb and showed a bimodal distribution (Supplementary Figure S5). The species *P. brasiliense* and *P. versatile* showed the most haplotypes, 21 and 27 respectively (Figure 2b). Five species had 8 or more TFL haplotypes and the remaining species had less than 5 haplotypes. This differential abundance of haplotypes in *Pectobacterium* species could be due to the sampling bias in our genome collection (See “genome count” in Figure 2b). Interestingly, intra-species average nucleotide identity (ANI) is not related to the number of TFL haplotypes. For example, *P. brasiliense* is more diverse than *P. versatile* (based on broader intra-species ANI in *P. brasiliense* (Supplementary Figure S6)), however, we observed more TFL haplotypes in the ecologically and geographically broadly distributed *P. versatile* (Portier et al. 2020) compared to *P. brasiliense*.

### The same tail fiber loci are observed across different species

We observed that genomes from two or more *Pectobacterium* species can have the same TFLs (Figure 2b, Supplementary Table S6). For example, among the samples collected at the watershed niche along the Durance River in the southeast of France (Ben Moussa et al. 2022), *P. carotovorum* (g_106, g_57) had the same TFL as *P. versatile* (g_53). In the Netherlands, we found that *P. brasiliense* (g_145, g_194, g_429) and *P. aroidearum* (g_421, g_442) have the same TFL haplotype. *P. versatile* (g_150, g_447) and *P. odoriferum* (g_434) from the Netherlands likewise shared the same TFL haplotype. In contrast, some species did not share TFL haplotypes, for example *P. parmentieri* and *P. brasiliense*. To further understand the complete extent of TFL haplotype sharing, we mapped all TFL haplotypes on the *Pectobacterium* genus phylogeny (Figure 3a-c). We observed that these haplotypes do not follow the phylogeny and found widespread evidence of shared TFL haplotypes, both intra- and inter-species.

**Figure 3:**
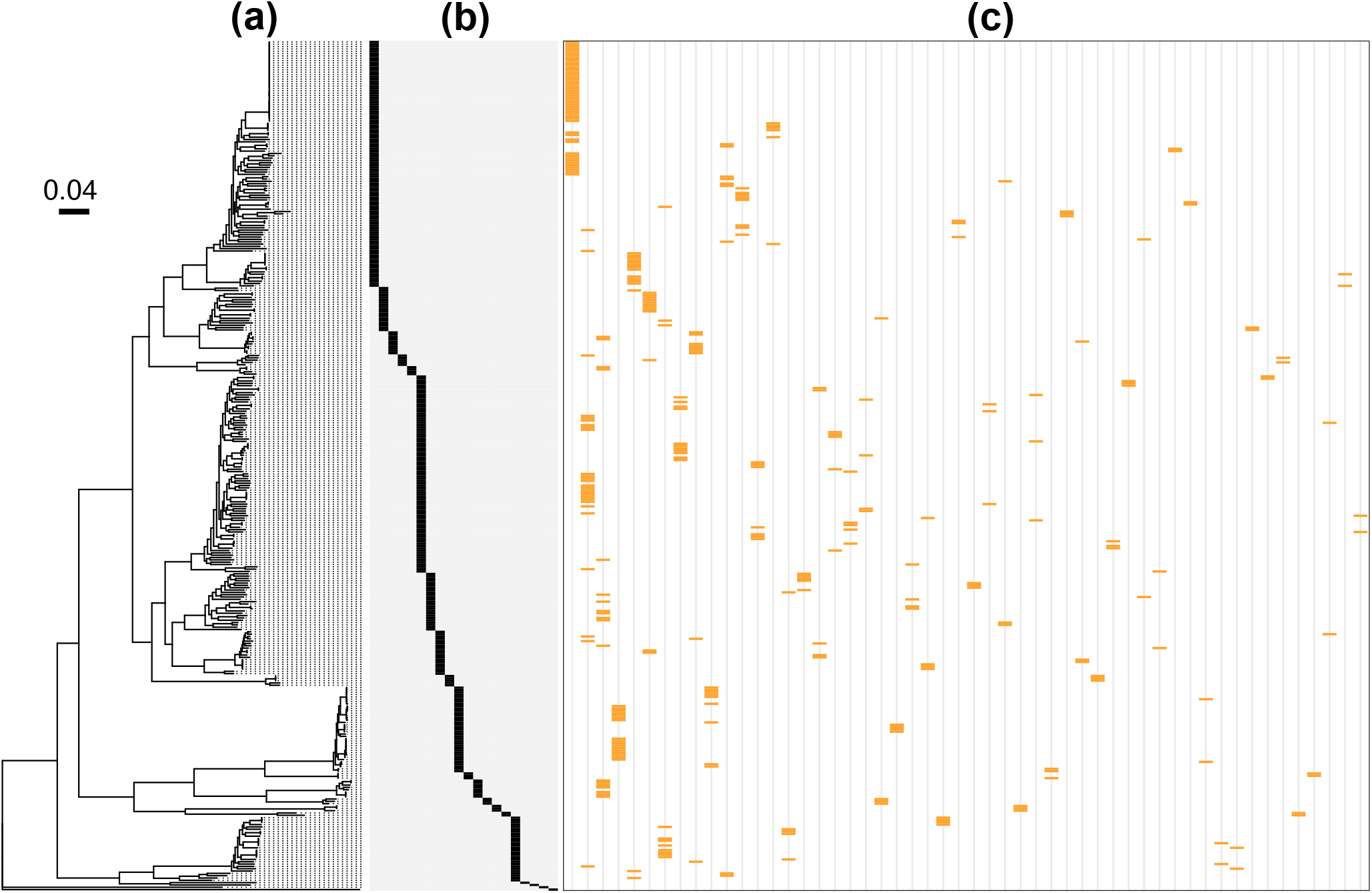
Carotovoricin polymorphism due to tail fiber locus diversity. (a) A core-genome phylogenetic tree is shown for the 365 *Pectobacterium* genomes that contain the carotovoricin cluster in a single contig. (b) A species key marking species from left to right: *P. brasiliense, P. polaris, P. parvum, P. aquaticum, P. quasiaquaticum, P. versatile, P. carotovorum, P. odoriferum, P. actinidiae, P. parmentieri, P. wasabiae, P. punjabense, P. polonicum, P. zantedeschiae, P. peruviense, P. aroidearum, Pectobacterium sp. CFBP8739, P. colocasium, P. fontis* and *P. cacticida*. (c) The presence of 52 TFL haplotypes is shown as yellow bands. Only the haplotypes that are present in two or more genomes (52 out of 93) are shown.

To study the intra- and inter-species variation of the conserved locus and the TFL (as defined in Figure 2a) at a higher resolution, we aimed to compare the DNA sequences between these two loci across the *Pectobacterium* genus. However, the presence of multiple structural variants in the TFL region, including inversions, insertions, and deletions, impedes the reconstruction of a reliable multiple sequence alignment, which would be required for phylogeny reconstruction. Therefore, we used a *k*-mer approach to calculate pairwise distance between these loci. Pairwise species distance was calculated as the cophenetic distance in the core-SNP phylogeny derived from the pangenome.

Under the assumption of uniform mutation rate across the genome, we expect the locus distance to correlate with the species distance. In line with this, pairwise locus distances for the conserved locus showed a strong positive correlation with species distance (*R* = 0.86, *p* < 2.2 × 10^−16^) (Figure 4a). Additionally, the intraspecies pairwise locus distances for the conserved locus were smaller than the inter-species distances, as might be expected. For the TFL, we also observed a positive, albeit weaker, correlation between locus distance and species distance (*R* = 0.55, *p* < 2.2 × 10^−16^) (Figure 4b). However, the locus distance range was significantly broader for both intra- and inter-species TFL compared to the conserved locus (Levene’s test, *p* < 2.2 × 10^−16^). This overall broad range of pairwise distances for TFLs underlines the sequence level diversity of the tail fiber pool in the *Pectobacterium* genus. As expected, TFL pairs with the same haplotypes exhibited lower locus distances, regardless of whether they were intra- or inter-species comparisons. Strikingly, however, some of the inter-species TFL pairs had a very small locus distance (sometimes even zero), indicating (nearly) identical TFLs in different *Pectobacterium* species (Figure 4b).

**Figure 4:**
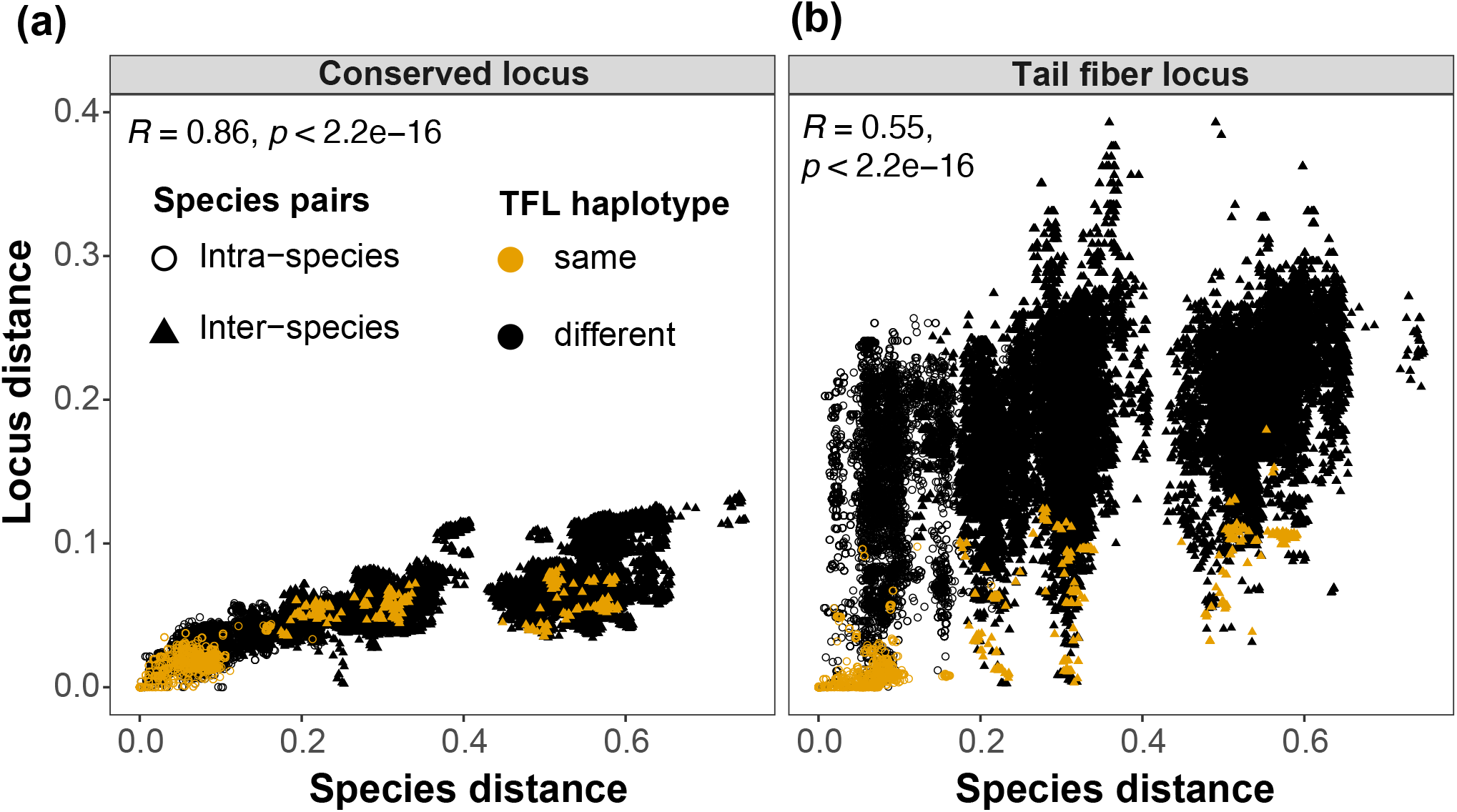
Sequence diversity of the tail fiber locus. Pairwise locus distances for carotovoricin regions, estimated using Mash, were compared against the species evolutionary distance, estimated from the *Pectobacterium* core-SNP phylogeny derived from the pangenome. These comparisons are shown for two loci in the carotovoricin: (a) the conserved locus, and (b) the variable TFL. The comparisons in which both loci belong to the same TFL haplotype are colored in yellow in both scatter plots for reference. Such matching haplotypes show smaller locus distances, also at higher species distances, for inter-species comparisons of TFLs (b). This trend is not observed for the conserved loci in (a).

### Tail fiber loci undergo genus-wide homologous recombination-mediated exchange

Given the presence of identical TFLs across different species, suggesting a possible horizontal gene transfer (HGT), we further analyzed phylogenetic incongruence between the conserved locus and the TFL using 16 genomes from 10 *Pectobacterium* species. These genomes encompassed three different TFL haplotypes that lacked inversions, thus allowing us to reconstruct a reliable multiple sequence alignment and phylogenetic tree. In the absence of HGT, the expectation is that both phylogenetic trees should show a consensus and follow the *Pectobacterium* species tree. However, while the conserved locus phylogeny showed strains clustering by *Pectobacterium* species, the tree derived from the variable TFL region alignment did not follow the *Pectobacterium* species phylogeny (Figure 5a-b). This therefore supports that the TFLs are exchanged among *Pectobacterium* species, independent of the rest of the carotovoricin locus.

**Figure 5:**
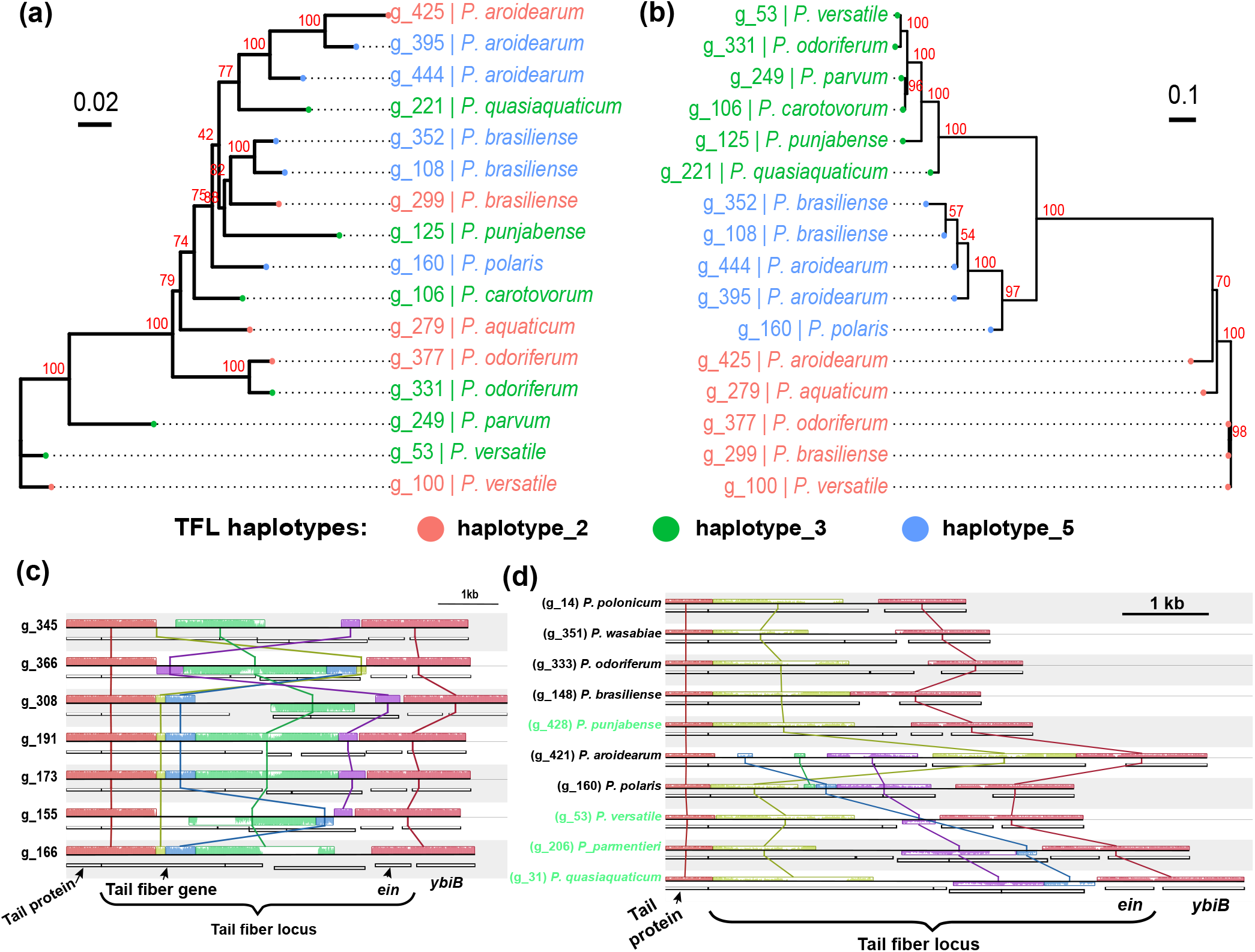
Horizontal gene transfer of tail fiber loci. (a) An unrooted maximum-likelihood phylogenetic tree for the conserved locus of carotovoricin from 16 genomes. (b) An unrooted maximum-likelihood phylogenetic tree generated from the TFLs from 16 genomes comprising 3 different haplotypes. For both the trees, bootstrap values are shown next to internal nodes and tips are colored as per the TFL haplotype. (c) Complex structural rearrangements of tail fiber locus (TFL) haplotypes are shown in the form of Mauve collinear blocks for carotovoricin loci in *P. brasiliense* accessions with the invertase gene *ein*. (d) Mauve collinear blocks in TFLs of various *Pectobacterium* species are shown and labels for TFLs that have invertase *ein* are colored green. Locally collinear blocks are shown as colored boxes above (forward) and below (reverse) the black horizontal line that represents the TFL haplotype region in a genome. Conserved blocks across the genomes are linked by vertical lines. Inside each block, a similarity profile is drawn which represents the average level of conservation across the TFL haplotypes. White areas represent sequences specific to a TFL and are not aligned.

A horizontal transfer of such loci between two species and strains can be mediated by homologous recombination. A key requirement for successful DNA recombination is the presence of a homologous region. To assess the existence of such compositional similarity at the sequence level in the regions flanking the TFLs, we identified locally collinear blocks of DNA sequences in selected TFLs using Mauve (A. C. Darling et al. 2004; A. E. Darling et al. 2010).

First, the intra-species comparison of *P. brasiliense* TFLs with *ein* showed inversions in this region across several genomes, flanked by positionally conserved locally collinear blocks with high conservation across the genomes (Figure 5c). These positionally conserved collinear blocks imply a common mechanism of DNA inversion, most likely Ein-mediated. The upstream conserved collinear block in TFL overlaps with the first half of the tail fiber gene, implying a conserved N-terminal domain of the protein encoded by this gene. This observation extends the previously reported modular structure of tail fibers in *P. carotovorum* (Yamada et al. 2006) to the other *Pectobacterium* species. The remaining region of the TFL shows complex structural rearrangements and results in variable C-terminal tail fiber domains.

Another comparison involving TFLs from multiple *Pectobacterium* species also showed a complex reordering and inversion of locally collinear blocks (Figure 5d). The flanking collinear blocks in TFLs without *ein* also showed positional conservation (Figure 5d), although it was less pronounced. The overall conservation of positions of the collinear blocks flanking the TFLs in both sets supports our hypothesis of recombination-mediated horizontal gene transfer.

## Discussion

Carotovoricin, a tailocin, is produced constitutively as well as under stress by *Pectobacterium* species during host infection, both on symptomatic and surrounding healthy tissue (Borowicz, Krzyżanowska, Sobolewska, et al. 2025). Previous studies have indicated that the tailocins target the strains of producer species or closely related species. Therefore, the target spectrum diversity of these tailocins plays a critical role in shaping the bacterial community (Heiman et al. 2023). The comparative pangenomics approach used in this study captured the diversity of carotovoricin in the *Pectobacterium* genus. We identified a variable degree of conservation within the carotovorocin cluster and a genus-wide horizontal gene transfer of tail fiber loci. Here we discuss the observed evolutionary dynamics and suggest new research directions based on this work.

### Evolutionary dynamics

In order to understand the scope of tailocin-mediated interactions in *Pectobacterium*, we conducted a genuswide analysis of their organization and evolution. Here we discuss the possible implications of the observed evolutionary dynamics in *Pectobacterium* and compare them with other tailocin-producing species.

Conservation of both carotovoricin core functional genes and their chromosomal location across the *Pectobacterium* genus, shown previously (Arizala and Arif 2019; Borowicz, Krzyżanowska, Sobolewska, et al. 2025; Pardeshi et al. 2025) and at a higher resolution in this study, underscores the important role of this tailocin in competitive interactions. The competitive interactions experienced by the bacteria may act as a negative selection pressure to preserve the tailocin functionality, evidenced by carotovoricin loss in less competitive and host-deprived niches (Supplementary Results and Discussion). This selection pressure is further emphasized by the observation that carotovoricin was found to be present in a single copy and no other tailocin was identified in *Pectobacterium* species genomes. This is different from *Pseudomonas* species, where multiple tailocins exist with various integration sites, evolutionary history and regulatory genes (Ghequire, Dillen, et al. 2015; Hockett et al. 2015). We observed species-wide partial degradation of carotovoricin cluster in *P. betavasculorum* and *P. atrosepticum*. Although we found transcriptional evidence for genes in the partially deleted carotovoricin cluster of *P. atrosepticum*, the function of such partial tailocin biosynthesis gene clusters remains unclear, as they do not code for a complete functional unit. Nonetheless, this indicates that the phage tail-like bacteriocins are not immune to dynamic degradation, a phenomenon observed in their prophage ancestors where a rapid initial deactivation is followed by gradual decay (Bobay et al. 2014). We hypothesize that their role in inter-bacterial interactions likely imposes a purifying selection pressure that protects tailocins from rapid initial deactivation, resulting in different domestication dynamics of tailocins compared to prophages.

Although carotovoricin is broadly conserved in the *Pectobacterium* genus, we observed varying degrees of conservation for multiple loci within the carotovoricin cluster, namely the nitric oxide deoxygenase gene, the tape measure gene and the tail fiber locus. We focused primarily on the TFLs because variation in their number and composition is a generic property of tailocins that directly determines the target specificity (Patz et al. 2019). Multiple sequence alignment has been used to estimate TFL haplotypes for *Pseudomonas* species tailocins (Backman, Latorre, et al. 2024). However, generating a multiple sequence alignment for genomic regions with complex structural variation, such as the TFLs, is challenging. Our pangenome approach to identify TFL haplotypes using homology group signatures overcomes this complexity.

We hypothesize that the *Pectobacterium* community in a plant host niche maintains a pool of carotovoricin tail fiber diversity to resist invasion by other closely related *Pectobacterium* species pathogens. Indeed, such a competitive exclusion mechanism is noted for R-tailocin producing *Pseudomonas chlororaphis* in mixed-species rhizosphere communities (Dorosky, Pierson, et al. 2018; Dorosky, Yu, et al. 2017). Additionally, the maintenance of TFL diversity in *Pectobacterium* might be over both space and time, as reported for another phytopathogen, *Pseudomonas* (Backman, Latorre, et al. 2024). Such a mechanism could be beneficial for the established colonizers in case of multi-species soft-rot infections, which are commonly observed in *Pectobacterium* species pathogens (Pardeshi et al. 2025; Smoktunowicz et al. 2022; Zhou et al. 2022).

Unexpectedly, the known mechanism to achieve TFL diversity, Ein-mediated inversion (Nguyen et al. 2001), was absent in approximately half of the carotovoricins within the *Pectobacterium* genus. However, we showed the existence of an additional mechanism to acquire the TFL diversity: horizontal transfer of TFLs across the *Pectobacterium* genus. Phylogenetic incongruence showed that HGT occurs in the region encoding the tail fiber of the carotovoricin and is likely mediated by site-specific recombination. Although the recombinational exchange of TFLs is known in other tailocin-producing bacteria (Baltrus, Clark, Smith, et al. 2019; Fautt et al. 2025), a combination of the two mechanisms, complementing each other, to maintain and circulate a pool of TFLs make *Pectobacterium* phytopathogens a unique case, to our knowledge. Additionally, it would be intriguing to test whether the HGT mechanism we found in the *Pectobacterium* genus can also be found in other genera involved in soft-rot plant diseases, for example, the *Dickeya* species.

Given the role played by carotovoricin in competitive interactions, it is puzzling why *P. atrosepticum* and *P. betavasculorum* never reacquired a functional copy of carotovoricin by HGT from other *Pectobacterium* species. One explanation could be that HGT of a complete carotovoricin locus is not possible due to its size or required homology, as we only observed transfer of TFLs in the *Pectobacterium* genus. Another possibility is that these species do not co-exist with other *Pectobacterium* species, because of niche adaptation or other inter-species antagonistic interactions (Smoktunowicz et al. 2022). Co-existence is essential for any successful HGT between two bacteria. Multiple *Pectobacterium* species share ecological niches and even cooccur on both symptomatic and asymptomatic plant material (Motyka-Pomagruk et al. 2021; Smoktunowicz et al. 2022; Zhou et al. 2022). In line with this, common TFL haplotypes are observed between such coexisting species. In contrast, species that do not co-exist, specifically *P. brasiliense* and *P. parmentieri* (Smoktunowicz et al. 2022), do not share any TFL haplotypes.

The success and stability of a heterogeneous community depend on having the means to overcome competition from other closely related strains or species. Thus, the ability to exchange TFLs adds a new dimension to the carotovoricin-mediated community dynamics involving multiple *Pectobacterium* species and strains. Based on the observed TFL diversity and the sharing of TFL haplotypes across species, innovations in terms of functional diversity appear to be shared within the *Pectobacterium* genus.

### New research directions

Our new insights on carotovoricin evolution in the *Pectobacterium* genus opens up various alleys for future data-driven research to study bacterial interactions. Here we discuss two promising directions.

First, our work provides a foundation for data-driven experimental design of tailocin competition assays to discover new interaction rules between multiple species. Recently, inter-genus killing activity of tailocin was shown in *Dickeya* species, which could target several *Pectobacterium* species (Borowicz, Krzyżanowska, Sobolewska, et al. 2025). Therefore, a data-first approach could be used to select the candidate strains across multiple species and spanning different TFL haplotypes to perform tailocin interaction experiments. Such an experimental design would facilitate the development of tailocin interaction prediction models, which can further resolve the tailocin-mediated interactions in bacteria. Due to carotovoricin cluster conservation, presence of a single copy per strain and localization at the same gene neighborhood across the genus, we propose to use *Pectobacterium* as a model to study inter-species tailocin mediated interactions.

Second, a fundamental understanding of tailocin diversity is essential to design novel tailocin-based applications targeting a specific group of pathogens. Recently, several of these applications have been developed, ranging from phytopathogen control (Baltrus, Clark, Hockett, et al. 2022; Ishii et al. 2024; Kouzai et al. 2024), engineering broad-range tailocins from phages (Williams et al. 2008; Woudstra et al. 2024) as well as designing broad-range chimeric phages based on the tailocins (G. Yao et al. 2023). The hyper-variable regions, such as the tail fiber loci, are difficult to compare at a large scale using a traditional comparative genomics approach. The pangenome approach taken here allows a detailed characterization of these TFLs and their contribution to tailocin evolution, which is essential for the development of tailocin-based biocontrol applications. With a growing number of high quality genomes in public databases, our comparative pangenomics approach provides a template to explore this as well as discover whether similar or novel mechanisms exist in other bacterial species.

## Materials and Methods

### Genome collection and pangenome construction

A *Pectobacterium* genus pangenome comprising 454 genomes was recently constructed by us (Pardeshi et al. 2025). Briefly, we used PanTools (v4.1.1) (Jonkheer et al. 2022; Sheikhizadeh et al. 2016) to build a De Bruijn graph pangenome, including gene annotations, functional annotations using InterProScan (v5.56-89.0) (Blum et al. 2021; Jones et al. 2014) and EggNOG-mapper (v2.1.10) (Cantalapiedra et al. 2021; Hernández-Plaza et al. 2023). We optimized the clustering parameters at which genes are grouped together in homology groups. For example, a strict setting (allowing less sequence diversity) can result in orthologous genes being distributed into multiple homology groups, consequently inflating the total number of homology groups in the pangenome. Conversely, a relaxed setting (allowing for more sequence diversity) would yield a smaller number of homology groups by incorrectly aggregating diverse genes into a single group. To study the carotovoricin evolutionary dynamics, the optimal grouping setting for the whole pangenome was estimated by evaluating the grouping of BUSCO gene sets under eight different settings (Jonkheer et al. 2022). The PanTools homology grouping setting of 4 (equivalent to >65% sequence identity, 0.05 intersection rate for the *k*-mer set comparison, a 7.2 inflation rate for Markov clustering and a contrast factor of 5) optimally organizes BUSCO gene sets in the *Pectobacterium* genus.

### Phylogenetic analysis

Multiple sequence alignment of genes in each of the 1,949 core homology groups was performed using the PanTools *msa* function. These alignments were used to construct a maximum likelihood phylogeny using the *core_phylogeny* function (Minh et al. 2020), which uses a concatenation-based phylogenomic approach. Phylogenetic trees were processed using R (v4.2.1) packages Ape (v5.7.1) (Paradis, Claude, et al. 2004; Paradis and Schliep 2019) and treeio (v1.22.0) (Wang et al. 2020) and visualized using ggtree (v3.6.2) (Xu et al. 2022) and ComplexHeatmap (v2.15.1) (Gu et al. 2016).

### Carotovoricin identification

Prophages were predicted using the geNomad (v1.5.0) pipeline (Camargo et al. 2023) and completeness was assessed using checkV (v1.0.1) (Nayfach et al. 2021). Any bacterial genomic regions at the prophage borders were identified using CheckV and subsequently trimmed. Putative prophages without any gene of viral origin were not considered. Each prophage was represented as a sequence of pangenomic homology groups, referred to as a prophage signature. When multiple prophage signatures from a genome could map in tandem to a larger prophage signature in another genome with *≥* 80% coverage, prophages from the first genome were considered as fragmented. Such fragmented prophages were merged into a single one and labeled as fragmented (110 fragmented prophages were merged into 50). All putative prophages with length < 5kb were further excluded, resulting in a final collection of 1,369 prophages.

The pairwise similarity between prophage homology group signatures was quantified as the syntenic conservation of homology groups: the syntenic Jaccard index. To calculate the syntenic Jaccard index, dynamic programming was used to align two prophage signatures, with a score of +5 for a match and −2 for a mismatch of homology groups between two prophage signatures. The longest common subsequence between two prophage signatures, with at most two gaps and at least five matching homology groups, was considered a valid alignment of homology groups. The syntenic matching homology groups from such valid alignments were used to determined the intersection size while calculating the Jaccard index. The syntenic Jaccard index matrix for all-vs-all comparisons was clustered using complete linkage hierarchical clustering with the ‘hclust’ function in R. Next to various small clusters, this yielded one large group of 428 orthologous prophages. Among these, prophages from *P. carotovorum* matched the phage tail-like bacteriocin encoding region, called “Carotovoricin Er”. All other prophages belonging to this cluster showed similar gene structures and hence were denoted as carotovoricin.

### Transcriptomics evidence for carotovoricin in *P. atrosepticum*

Normalized count data for wild-type and polyphenol treated RNAseq samples for *P. atrosepticum* was downloaded from the GEO record GSE196675 (Kang et al. 2022). Genes from the partial carotovoricin locus were located in the region NC_004547.2:3113731-3119520. These genes were found to be differentially expressed upon polyphenol treatment.

### Carotovoricin region synteny visualization

A 5 kb region flanking the carotovoricin cluster was included to provide broader context to understand integration at the same or different site in the genome. An R script was written to extract the homology groups for carotovoricin regions and the data was stored in the JSON format required by clustermap.js (https://github.com/gamcil/clustermap.js). For species wise visualization for *P. brasiliense* and *P. versatile*, carotovoricins for the respective species were clustered based on the homology group presenceabsence matrix and the dendrogram was cut at the appropriate height to select representatives for clinker plotting.

### Comparative phylogenetics analysis of tail fiber locus

The carotovoricin region from the tail protein (hg_22427604) to *ybiB* (hg_22427603) was defined as the tail fiber locus (TFL) and the remaining upstream region as the conserved locus. Haplotypes for the tail fiber locus were determined based on the unique ordering of homology groups in this region across all *Pectobacterium* genomes. Locus distance, estimated as a sequence-based distance between carotovoricin pairs, was calculated using the MinHash approach in Mash (v2.3) (Ondov et al. 2016). Pairwise locus distance was calculated separately for the conserved locus and tail fiber locus. Pairwise species distances were calculated as the cophenetic distance using ‘cophenetic.phylo()’ function from the R package Ape (Paradis, Claude, et al. 2004; Paradis and Schliep 2019). Cophenetic distances are calculated as pairwise distances between the pairs of tips of a phylogenetic tree using its branch lengths.

Carotovoricins from 16 genomes spanning three TFL haplotypes and representing 10 *Pectobacterium* species were selected for performing comparative phylogenetics analysis. The DNA sequences for conserved loci and TFLs for these carotovoricin regions were independently aligned using MAFFT (v7.508) (Katoh et al. 2019). Maximum-likelihood phylogenetic trees for these two loci were estimated using IQ-TREE2 (2.2.0.3) (Minh et al. 2020). The trees were visualized using R (v4.2.1) package ggtree (v3.6.2) (Xu et al. 2022).

### Visualization of structural variation in tail fiber locus

DNA sequences of carotovoricin regions from the tail protein (hg_22427604) to *ybiB* (hg_22427603) were extracted as TFL sequences. The homology group level annotations from the PanTools’ pangenome and DNA sequences were combined into a GenBank file format using the *seqret* tool from the EMBOSS software suite (v6.6.0.0) (Rice et al. 2000). These GenBank format sequences were aligned and visualized using the ‘progressiveMauve’ command of the Mauve v(2.4.0) program (A. C. Darling et al. 2004; A. E. Darling et al. 2010).

## Supporting information

Supplementary results and figures

## Acknowledgements

We would like to acknowledge Inge van Duivenbode from Dutch General Inspection Service for agricultural seeds and seed potatoes (NAK) and Michiel J. C. Pel from the Netherlands Institute for Vectors, Invasive plants and Plant health (NIVIP) for providing *Pectobacterium* isolates and their valuable discussion in this research.

## Funding

This research was funded by the Dutch Ministry of Economic Affairs in the Topsector Program “Horticulture and Starting Materials” under the theme “Plant Health” (project number: LWV20.235) and its partners (NVWA, NAK, Naktuinbouw and BKD).

## Authors’ contributions

LP, SS, TL, DR: Conceptualization. LP: Data curation, Formal analysis, Investigation, Software, Writing – original draft. SS, TL, DR: Supervision, Funding acquisition, Project administration. LP, SS, TL, DR, AK: Writing - review and editing. All authors reviewed the manuscript.

## Conflicts of interest

The authors declare no competing interests.

## Supplementary data

**Supplementary tables:** Supplementary_Tables.xlsx

**Supplementary results and figures:** Supplementary_Material.pdf

## Data Availability

Genome assemblies used in this study were downloaded from the NCBI database. Assembly accessions and metadata for 454 genomes are listed in Supplementary Table S1. All scripts used for preprocessing, pangenome construction and analysis are available in the GitHub repository https://lakhanp1.github. io/Pectobacterium_pangenome. Notebooks to reproduce the analysis, tables and figures can be accessed at https://lakhanp1.github.io/Pectobacterium_pangenome/scripts/notebooks/carotovoricin_analysis.html. GitHub repository is archived in Zenodo and can be accessed using DOI: https://doi.org/10.5281/zenodo.16585541.

## Notes

https://lakhanp1.github.io/Pectobacterium_pangenome/scripts/notebooks/carotovoricin_analysis.html

### Competing Interest Statement

The authors have declared no competing interest.

### Summary of Updates

- Updated Results text to provide more clarity - Updated Methods to provide additional clarity on homology grouping - New Supplementary Figure S5

