## Supplementary results and figures for "Genus-wide homologous recombination of tail fibers maintains tailocin diversity in *Pectobacterium*"

### Supplementary data, belonging to Pardeshi ***et al.***, "Genus-wide homologous recombination of tail fibers maintains tailocin diversity in ***Pectobacterium***"

#### Supplementary Results and Discussion

##### Rare events of carotovoricin disruption

In addition to the species-wide loss of carotovoricin, there were seven strains from various *Pectobacterium* species that showed disruption of the carotovoricin cluster. One *P. brasiliense* isolated from potato plant (g_149) showed an insertion of a bacteriophage genome within the carotovoricin cluster. One *P. versatile* isolated from the potato plant (g_313) showed a deletion of five genes in carotovoricin: tail sheath, tail core, ferredoxin I, tail assembly chaperone and a partial tail tape measure gene. One *P. carotovorum* isolate (g_15) sampled from the Japanese angelica tree, recently reclassified as *P. araliae* (Sawada et al. 2024), showed complete loss of the carotovoricin locus. Finally, we identified four *P. brasiliense* isolates lacking carotovoricin, all sampled in Europe after 2017. A homology group signature comparison of these isolates showed that the flanking gene neighborhood was intact, ruling out assembly artifacts (Supplementary Figure S2). These four isolates were sampled from fresh water in the Netherlands and France (g_182, g_185 and g_236), which has shown to be a rare ecological niche for *P. brasiliense* (Ben Moussa et al. 2022), and from insect traps (g_177). Based on the core gene phylogenetic tree, these four isolates formed a monophyletic, yet non-clonal group (Supplementary Figure S1).

We also assessed whether other important systems involved in interaction are lost in these genomes. A detailed comparison with the other 134 *P. brasiliense* genomes showed that the type III secretion system (T3SS), involved in the injection of bacterial effector proteins into the eukaryotic host cells to modulate host response during infection (Costa et al. 2015; Green and Mecsas 2016), was also missing in these four isolates (Supplementary Figure S3). With the existing data, it is difficult to ascertain either independence or association in the deletion of these two systems. Notably, the type 2 secretion system (T2SS) and type 6 secretion system (T6SS) remained intact in these four *P. brasiliense* genomes. The aquatic niches from which these strains were isolated are characterized by lower bacterial density and a lack of plant hosts, offering a different competition profile than agricultural niche. Given that tailocin production is dependent on environmental cues, with its induction occurring in the infected plant host but absent in nutrient-deficient environments such as river water (Borowicz et al. 2025), a strictly aquatic niche may not exert strong purifying selection pressure to maintain a function that is not used. Therefore, we hypothesize that prolonged growth of these *P. brasiliense* strains in less competitive and host-deprived niches may have resulted in the loss of both carotovoricin and T3SS.

#### Supplementary Figures

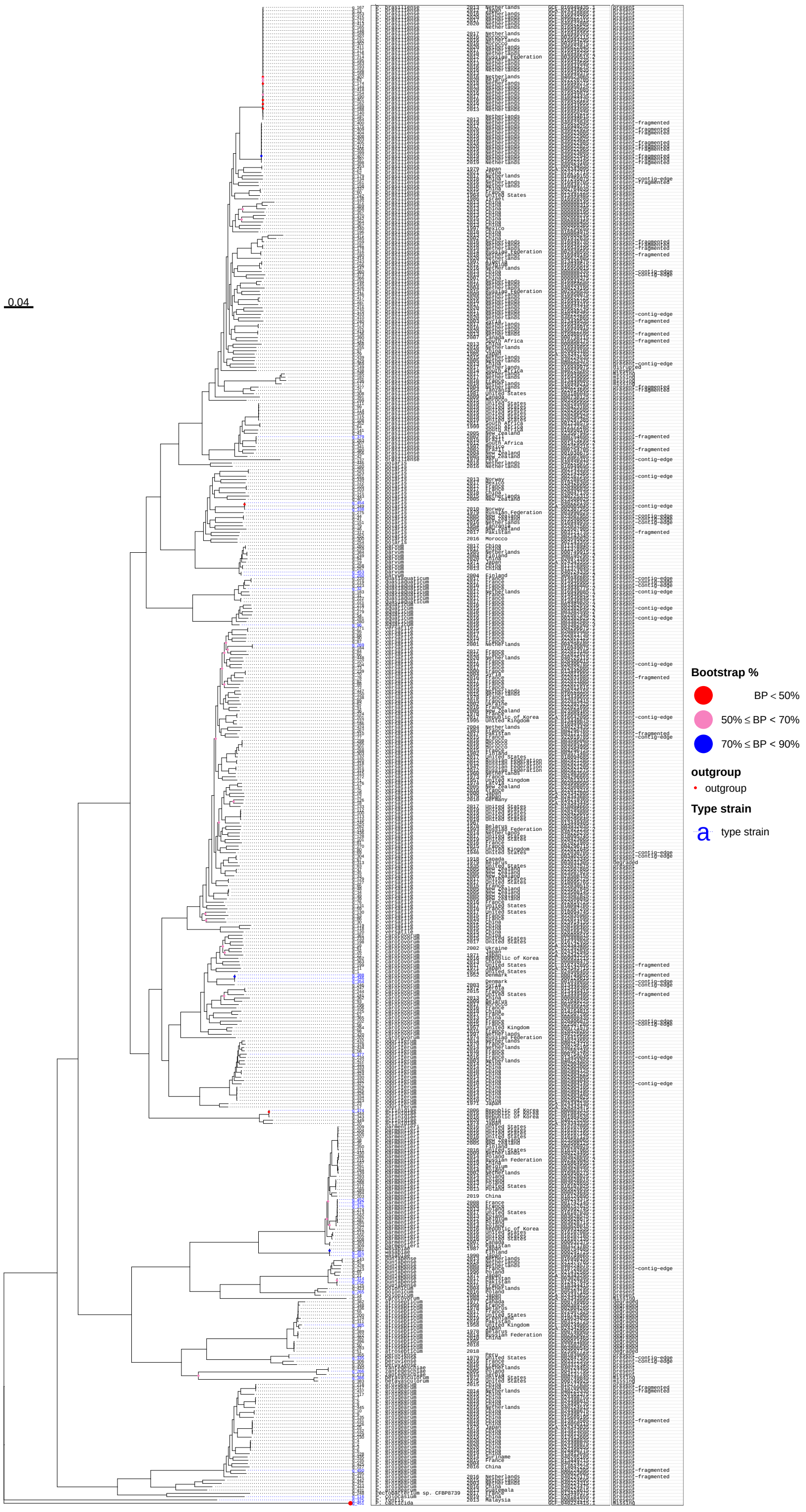


**Supplementary Figure S1:** *Pectobacterium* genus phylogeny. Core-genome phylogenetic tree for *Pectobacterium* genomes used to construct the pangenome. The tree is rooted using *P. cacticida* as the outgroup. The tip labels include the following data: genome identifier in the pangenome, *Pectobacterium* species name, year of isolation, country of origin, NCBI assembly identifier and carotovoricin presence-absence status.

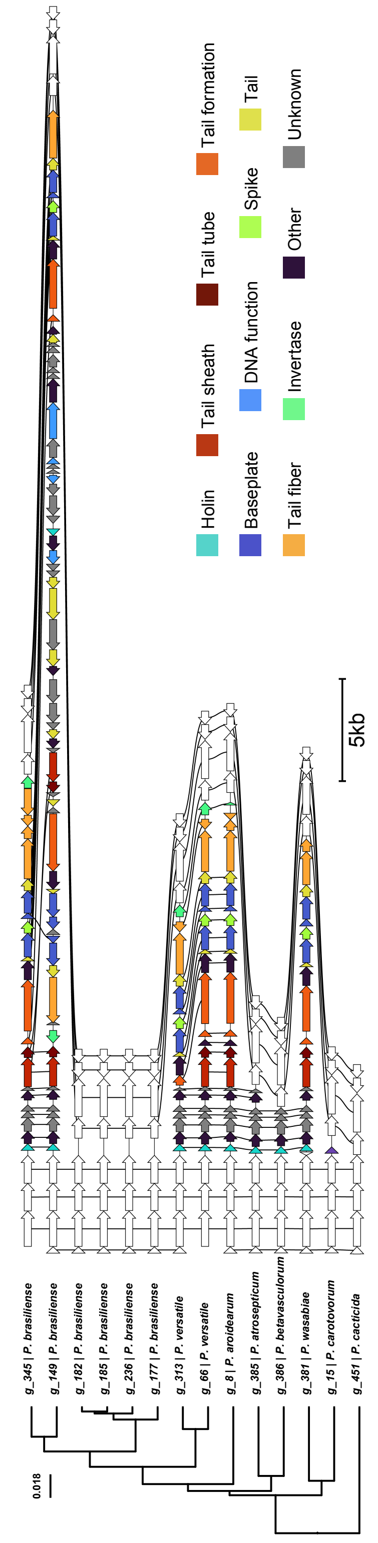


**Supplementary Figure S2:** Carotovoricin deletion. Genomic regions for strains in which carotovoricin was degraded are shown along with the phylogenetic tree. Genes in 5kb regions flanking carotovoricin clusters are shown in white. Genes belonging to the same homology group are connected with vertical black lines.

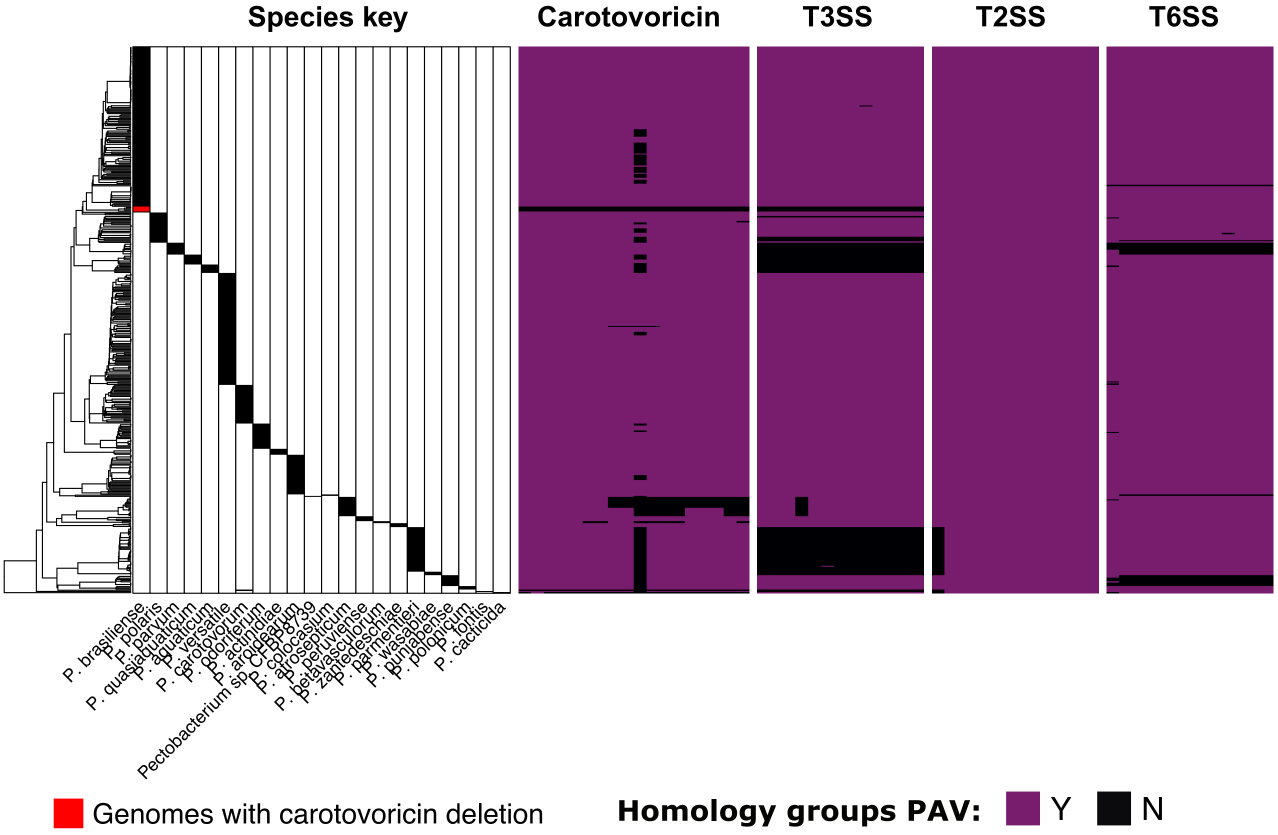


**Supplementary Figure S3:** Carotovoricin and T3SS deletion. *Pectobacterium* phylogeny for 454 genomes with a species key marking respective species column is shown on the left. *P. brasiliense* genomes with carotovoricin deletion are marked with the red color. The subsequent four heatmaps show core genes of carotovoricin, T3SS, T2SS and T6SS. In contrast to the T3SS genes, the T2SS and T6SS genes remain intact in the four *P. brasiliense* genomes.

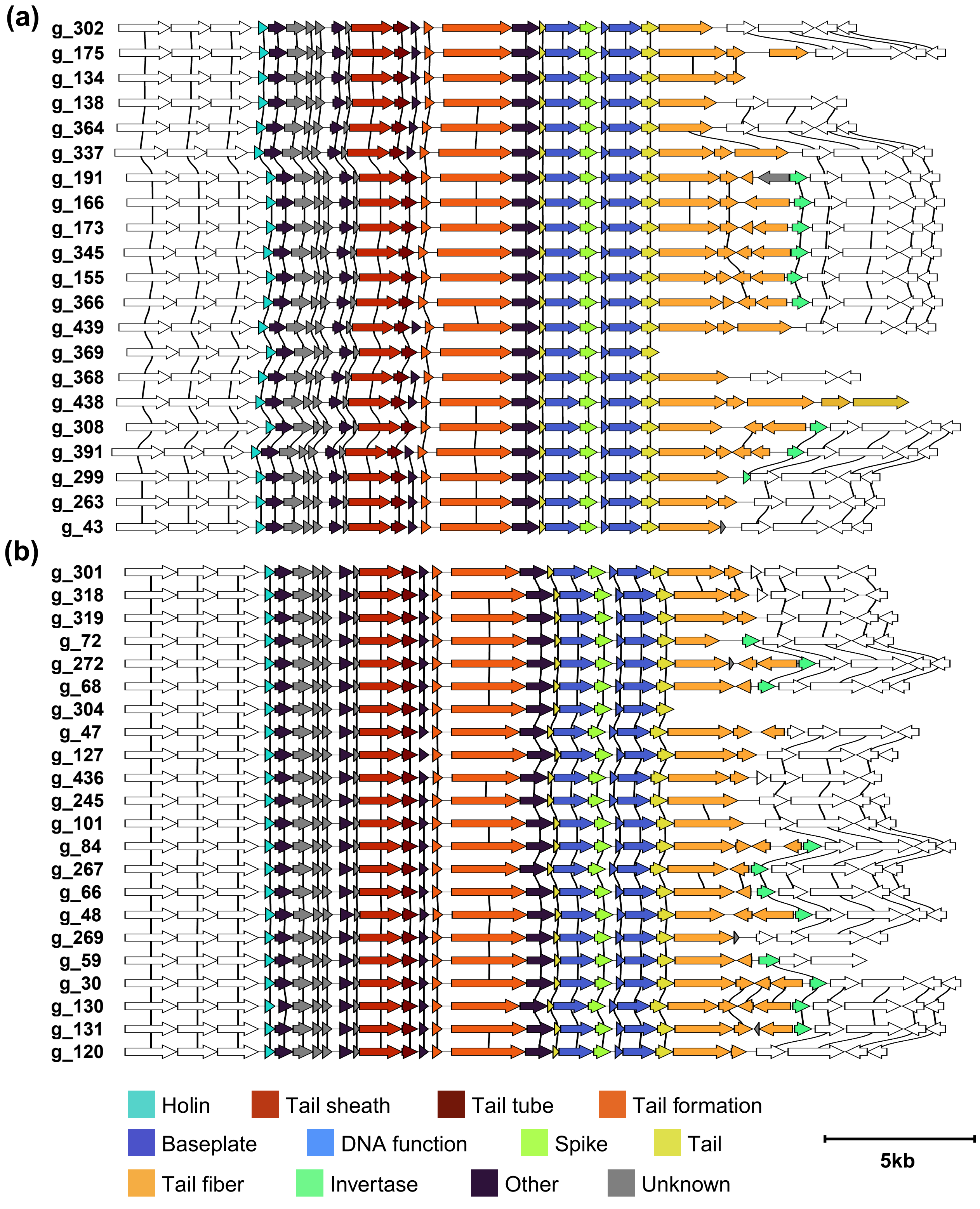


**Supplementary Figure S4:** Carotovoricin cluster variation in *Pectobacterium* species. Carotovoricin cluster is shown for the (a) *P. brasiliense* and (b) *P. versatile* species genomes. Genes in 5kb regions flanking carotovoricin clusters are shown in white. Genes belonging to the same homology group are connected with vertical black lines.

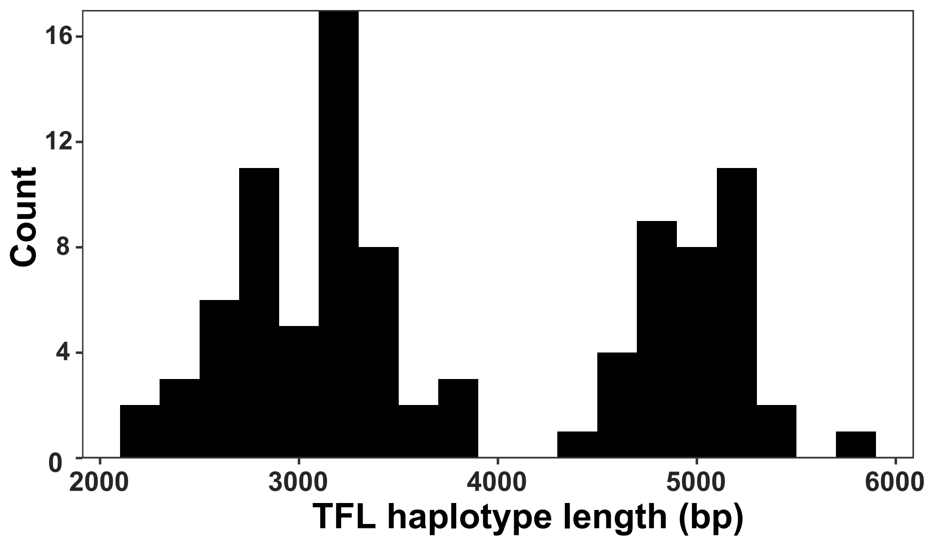


**Supplementary Figure S5:** Histogram of TFL haplotype length. The haplotype length is defined as the sequence length of the region after the tail protein up to the *ybiB* gene in the carotovoricin cluster.

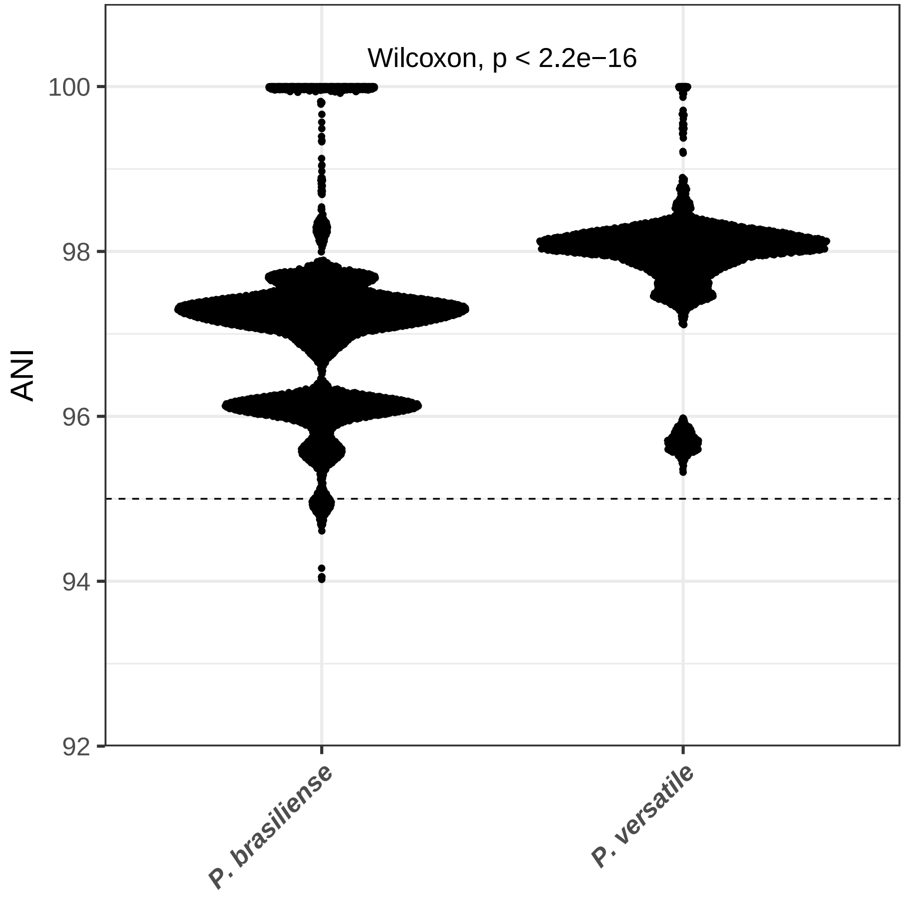


**Supplementary Figure S6:** Pairwise intra-species ANI comparison for *P. brasiliense* and *P. versatile* genomes. The mean values of intra-species ANI for these species are 97.1% and 97.9% respectively.
